# A glycerol shunt functions as a glucose excess security valve in pancreatic β-cells

**DOI:** 10.1101/2021.11.20.468884

**Authors:** Anfal Al-Mass, Pegah Poursharifi, Marie-Line Peyot, Roxane Lussier, Emily Levens, Julian Guida, Yves Mugabo, Elite Possik, Rasheed Ahmad, Fahd Al-Mulla, Robert Sladek, S.R.Murthy Madiraju, Marc Prentki

## Abstract

The recently identified glycerol-3-phosphate (Gro3P) phosphatase (G3PP) in mammalian cells, encoded by the *PGP* gene, was shown to regulate intermediary metabolism by hydrolyzing Gro3P and to control glucose-stimulated insulin secretion (GSIS) in β-cells, *in vitro*. We now examined in inducible β-cell specific G3PP-KO (BKO) mice, the role of G3PP in the control of insulin secretion *in vivo*, β-cell function and glucotoxicity. BKO mice, compared to *MCre* controls, showed increased body weight, adiposity, fed insulinemia, GSIS, reduced plasma triglycerides and mildly altered glucose tolerance. Isolated BKO mouse islets at high (16.7 mM) but not low or intermediate glucose (3-8 mM) showed elevated GSIS, Gro3P, metabolites reflecting β-cell activation, O_2_ consumption, ATP production and reduced glycerol release. BKO islets chronically exposed to elevated glucose showed increased apoptosis, reduced insulin content and expression of *Pdx-1, MafA* and *Ins-2* genes. As G3PP channels glucose carbons towards glycerol formation and release, the results demonstrate that β-cell are endowed with a “*glycerol shunt*” acting as a glucose excess security valve. We propose that the glycerol shunt plays a role in glucodetoxification, the prevention of insulin hypersecretion, acts as a defense against excess body weight gain and contributes to β-cell mass preservation in the face of hyperglycemia.

## Introduction

Hyperinsulinemia and insulin resistance contribute to obesity, type 2 diabetes (T2D) and associated metabolic disorders [1, 2]. Pancreatic β-cells secrete insulin primarily in response to increasing blood glucose levels, through the production of metabolic coupling factors (MCF) that promote insulin granule exocytosis. Disturbed glucose and lipid metabolism, including the glycerolipid/ free fatty acid (GL/FFA) cycle [3-5], which is implicated in the regulation of insulin secretion and energy homeostasis [6-8], contribute to the pathogenesis of obesity and T2D. The GL/FFA cycle, known to produce several MCF, in particular monoacylglycerol [9], consists of two arms: lipogenesis implicated in the synthesis of glycerolipids, and lipolysis during which glycerolipids, in particular triglycerides (TG) are hydrolyzed to FFA and glycerol [4, 5]. Lipogenesis involves the esterification of fatty acyl-CoA with glycerol-3-phosphate (Gro3P), which in β-cells is formed mostly from glucose via glycolysis. The GL/FFA cycle is driven largely by the availability of Gro3P and fatty acyl-CoA [3-5].

We recently reported that mammalian cells harbor a Gro3P phosphatase (G3PP), which by hydrolyzing Gro3P to glycerol, controls the availability of Gro3P for various metabolic pathways, in particular TG synthesis (lipogenesis segment of GL/FFA cycle), the Gro3P-electron shuttle to mitochondria and flux through lower glycolysis [10]. G3PP in mammalian cells, previously known as phosphoglycolate phosphatase, is encoded by *PGP* and we demonstrated that it primarily uses Gro3P as its intracellular substrate under normal physiological conditions. However, this enzyme is also able to use other phosphate esters, including phosphoglycolate as substrates, under stress conditions [11-13]. Our earlier *in vitro* studies primarily in INS1-ß-cells showed that G3PP activity is inversely proportional to glucose-stimulated insulin secretion (GSIS) at elevated glucose concentrations. We found that increased expression of G3PP in β-cells diverts a significant part of the glucose carbons towards the formation of glycerol, which is not metabolized due to low glycerol kinase activity [4, 14] and leaves the β-cell. Thus, elevated G3PP in β-cells prevents hypersecretion of insulin under excess glucose conditions by lowering the flux through the GL/FFA cycle, lower-glycolysis and mitochondrial oxidation (which are important sources of MCF), while the reverse was noticed with the suppression of G3PP [10]. In addition, glucotoxicity and glucolipotoxicity in β-cells were aggravated when G3PP expression was suppressed, whereas the cells were protected when G3PP was over-expressed, suggesting an important role of G3PP in the detoxification of glucose and protection of β-cells under nutrient-excess conditions [10].

However, the role of ß-cell G3PP in the control of insulin secretion and energy homeostasis *in vivo* is not known, nor do we know the range of glucose concentrations where the enzyme is active. In addition, if we have learned much about the pathways and biochemical basis of glucotoxicity, little is known as to how mammalian cells cope, adapt and detoxify excess glucose in the body [15]. Deletion of G3PP activity in the whole body was found to be embryonically lethal [12], and therefore we have generated a β-cell specific G3PP knockout (BKO) mice in which deletion of *PGP*/G3PP gene is induced in the adult stage, in order to avoid any developmental effects. The present *in vivo* and *ex vivo* studies, in which islets were exposed to low, intermediate and high glucose reveal that G3PP acts as a glucose excess security valve and identify a novel pathway of glucodetoxification, that we propose to name the “glycerol shunt”.

## Results

In the present study, we employed MCre mice as the control group for the G3PP-BKO mice considering that the MCre mice express Cre recombinase in response to tamoxifen, unlike the fl/fl and WT mice. Additionally, MCre mice were previously shown to display expression of human growth hormone sequences [16], which may influence islet function. Thus, it was previously suggested that for β-cell specific gene knockout studies using Mip-Cre-ERT mice [17], the better controls are the Mip-Cre-ERT mice than the WT or flox/flox (for the concerned gene) mice [18]. However, we compared the different control groups and found that WT and fl/fl mice have similar phenotype with regard to *in vivo* (Supplemental Figure 1) and *ex vivo* (Supplemental Figures 2 and 4) parameters. This justifies the use of MCre mice as control mice in this study.

### G3PP-BKO mice show increased body weight gain, fat mass and fed insulinemia

G3PP-BKO mice fed a standard chow diet till 20 weeks of age showed significant increase in body weight gain (Figure 1*D* and *E*) and fat mass (Figure 1*F*) with no change in food intake (Figure 1*G*) compared to MCre control mice. Even though there was no change in fed glycemia, fed insulinemia was significantly higher in the BKO mice with a similar tendency (*p* = 0.1) in C-peptide levels (Figure 1*H*). However, BKO mice showed lower plasma TG level with no changes in glycerol and FFA (Figure 1*H*). In accordance with the increased fat mass, we found that BKO mice have larger visceral adipose depots (Figure 1*I*) and higher visceral adipose tissue weight (Supplemental Figure 1*E*). There was no significant change in other tissue weights in BKO mice compared to controls and also among the different controls (Supplemental Figure 1*E*). The TG content of visceral adipose tissue and skeletal muscle was higher in the BKO mice (Figure 1*J*).

**Figure 1.**
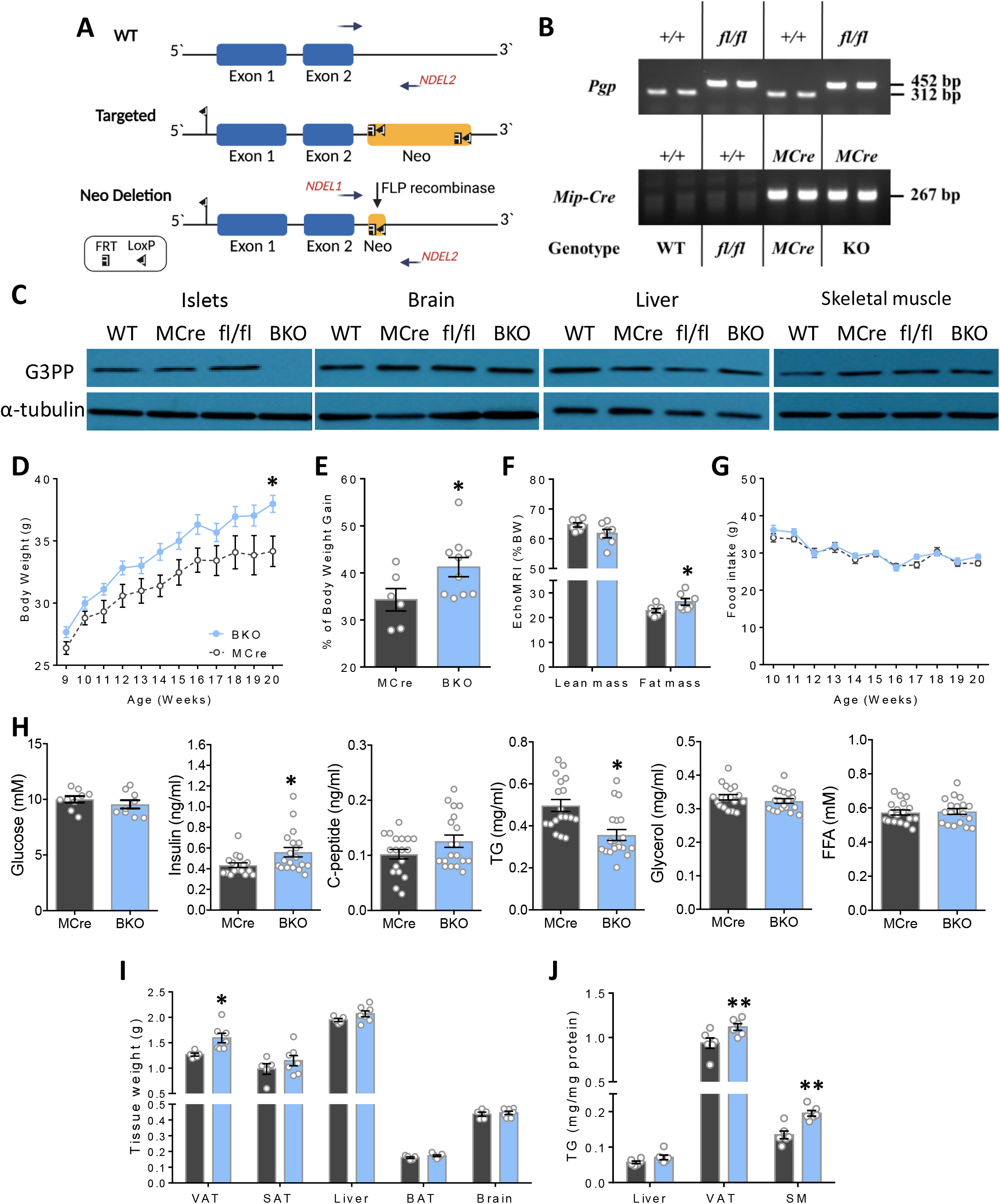
Increased body weight gain, fat mass and fed insulinemia with reduction in fed plasma TG in male BKO mice. (A) Schematic representation of gene targeting strategy for the generation of the G3PP-lox conditional allele. Wild-type, targeted and G3PP-lox alleles are represented. The loxP recombination sites inserted at the 5’ side of exon 1 and 3’ side of exon 2 in PGP gene, the FRP-flanked Neo-cassette FRP sites, and the NDEL1/NDEL2 primers for PCR genotyping are indicated. The breeding of mice to produce BKO mice is described in Methods. (B) The presence of the WT, *Mip-Cre* transgene or the floxed *PGP* alleles was evaluated in DNA from ear punch tissue fragments using the primers described in Methods. A specific amplification of a 452 bp DNA fragment corresponds to the floxed-*PGP* allele, a 312 bp fragment corresponds to the WT *PGP* allele and the presence of a 267 bp DNA fragment corresponds to the Mip-*Cre* transgene, as verified by genomic PCR. (C) G3PP deletion was validated using protein extracts from islets, brain, liver and skeletal muscle (SM) from mice, 4 weeks after deletion and used for western blot (WT, n=4; fl/fl, n=3; MCre, n=3; BKO, n=3). For all tissues α-tubulin was used as a housekeeping protein. Mice were kept on chow diet for 12 weeks after G3PP deletion following TMX treatment. (D) Body weight (Mcre, n=6; BKO, n=10). (E) Percentage of body weight gain (MCre, n=6; BKO, n=10). (F) Lean and fat mass expressed as percentage of BW (MCre, n=7; BKO, n=7 (one BKO mouse out of 8, died during experiment)). (G) Food intake (MCre, n=6; BKO, n=10). (H) Plasma parameters in fed state (For glycemia: MCre, n= 9 and BKO=9; for insulin, C-peptide, TG, glycerol and FFA, MCre, n=18; BKO, n=19). (I) Weights of visceral adipose tissue (VAT), subcutaneous adipose tissue (SAT), liver, brown adipose tissue (BAT) and brain (MCre, n=6; BKO, n=7). (J) TG content for liver, VAT and skeletal muscle (SM) (MCre, n=6; BKO, n=7). Data are mean ± SEM. *p < 0.05, **p < 0.01 vs MCre (Two-way ANOVA (Panel D and G) and Student’s *t* test (Panel E, F, H, I and J)).

### G3PP-BKO mice show increased insulin secretion *in vivo* and mild glucose intolerance

As the BKO mice displayed elevated fed insulinemia, we assessed glucose tolerance in these mice. Glucose tolerance, assessed by IPGTT, in the BKO mice was found to be modestly impaired, as indicated by the significantly elevated plasma glucose levels only at 30 min after intraperitoneal glucose injection, with larger AUC-glucose at 60 min, although no changes were seen at 15 min after the glucose load, compared to MCre controls (Figure 2*A*). Furthermore, plasma insulin levels were higher in the BKO mice at all the time points following glucose injection compared to control mice (Figure 2*B*). These results indicated that BKO mice secrete more insulin than control mice under similar hyperglycemic conditions. The IPITT suggested that the BKO mice fed a chow diet have unaltered insulin sensitivity (Figure 2*C*), despite showing elevated insulin secretion in response to a glucose load.

**Figure 2.**
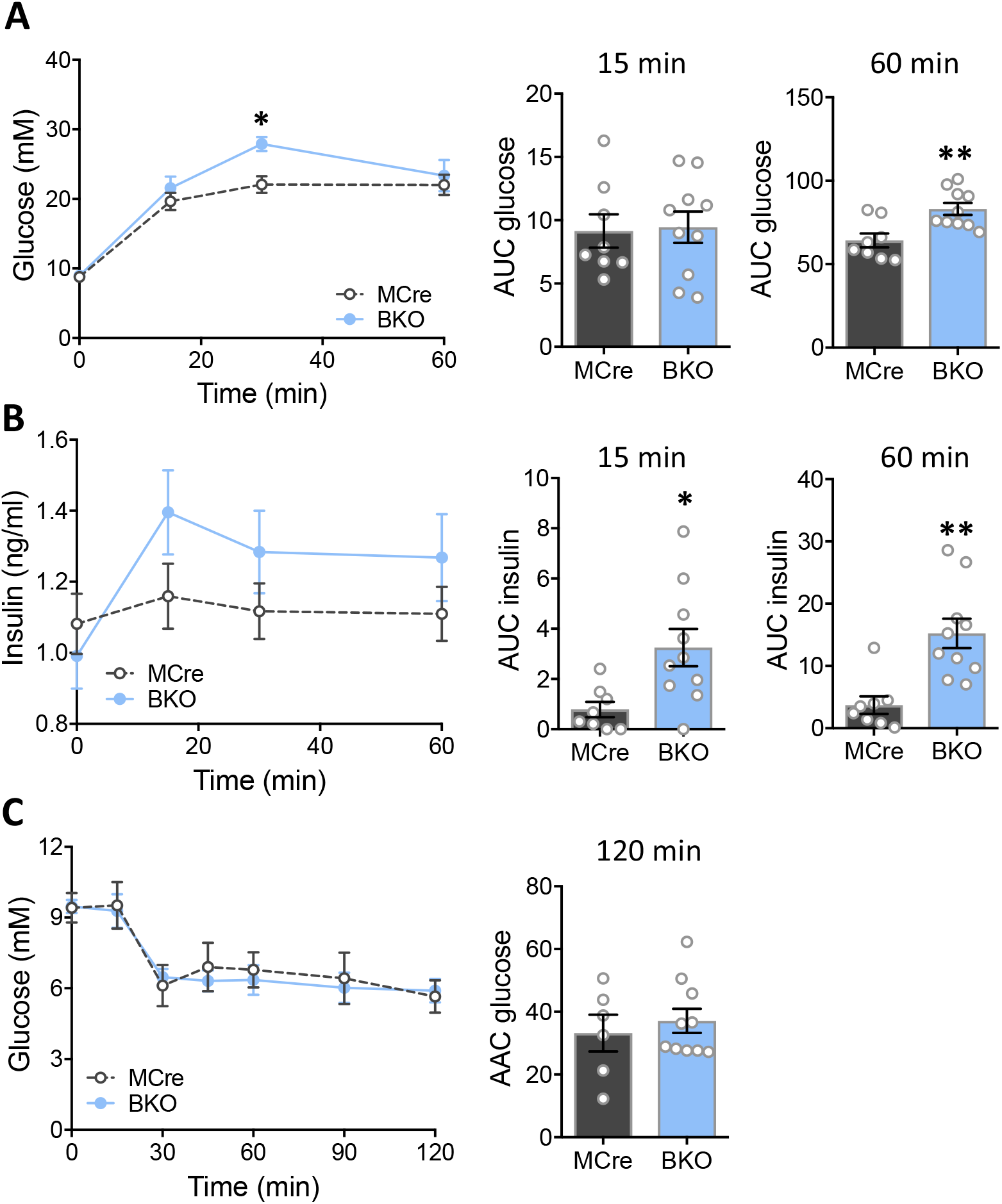
Increased *in vivo* glucose induced insulin secretion with mild glucose intolerance and normal insulin sensitivity in BKO mice. (A) Glycemia during IPGTT in male mice, 16 weeks after G3PP deletion (MCre, n=8; BKO, n=10). Inset depicts area under the curve (AUC) for glycemia after 15 min and 60 min. (B) Insulinemia during IPGTT in male mice (MCre, n=8; BKO, n=10). Inset depicts AUC for insulinemia after 15 min and 60 min. (C) Glycemia during ITT in mice, 19 weeks after G3PP deletion (MCre, n=6; BKO, n=10). Inset depicts area above the curve (AAC) after 120 min. Data are means ± SEM. *p < 0.05, **p < 0.01 vs MCre (Two-way ANOVA and Student’s *t* test (for AUC and AAC)).

### G3PP-BKO islets show increased insulin secretion and mitochondrial metabolism only at high glucose concentration

In order to examine if the *in vivo* deletion of *PGP* gene in mice impacts β-cell function, and if so in which glucose concentration range, we measured insulin secretion in BKO and control mouse islets *ex-vivo*, in response to different concentration of glucose with and without FFA. Insulin secretion at 3 mM (basal, hypoglycemia) and 8 mM (intermediate/physiological in mice) glucose concentration was similar between BKO and MCre control islets (Figure 3*A*). However, at high glucose concentration (16 mM) BKO islets showed markedly increased insulin secretion compared to the control islets (Figure 3*A*), indicating that G3PP activity is more relevant in the control of GSIS under hyperglycemic conditions. Similar results were obtained in the presence of FFA (Figure 3*B*). KCl-induced insulin secretion showed only a tendency to be higher in the BKO islets (Figure 3*C*). G3PP deletion in β-cells had no effect on insulin content (Figure 3*D*), but markedly lowered glucose-dependent glycerol release (Figure 3*E*).

**Figure 3.**
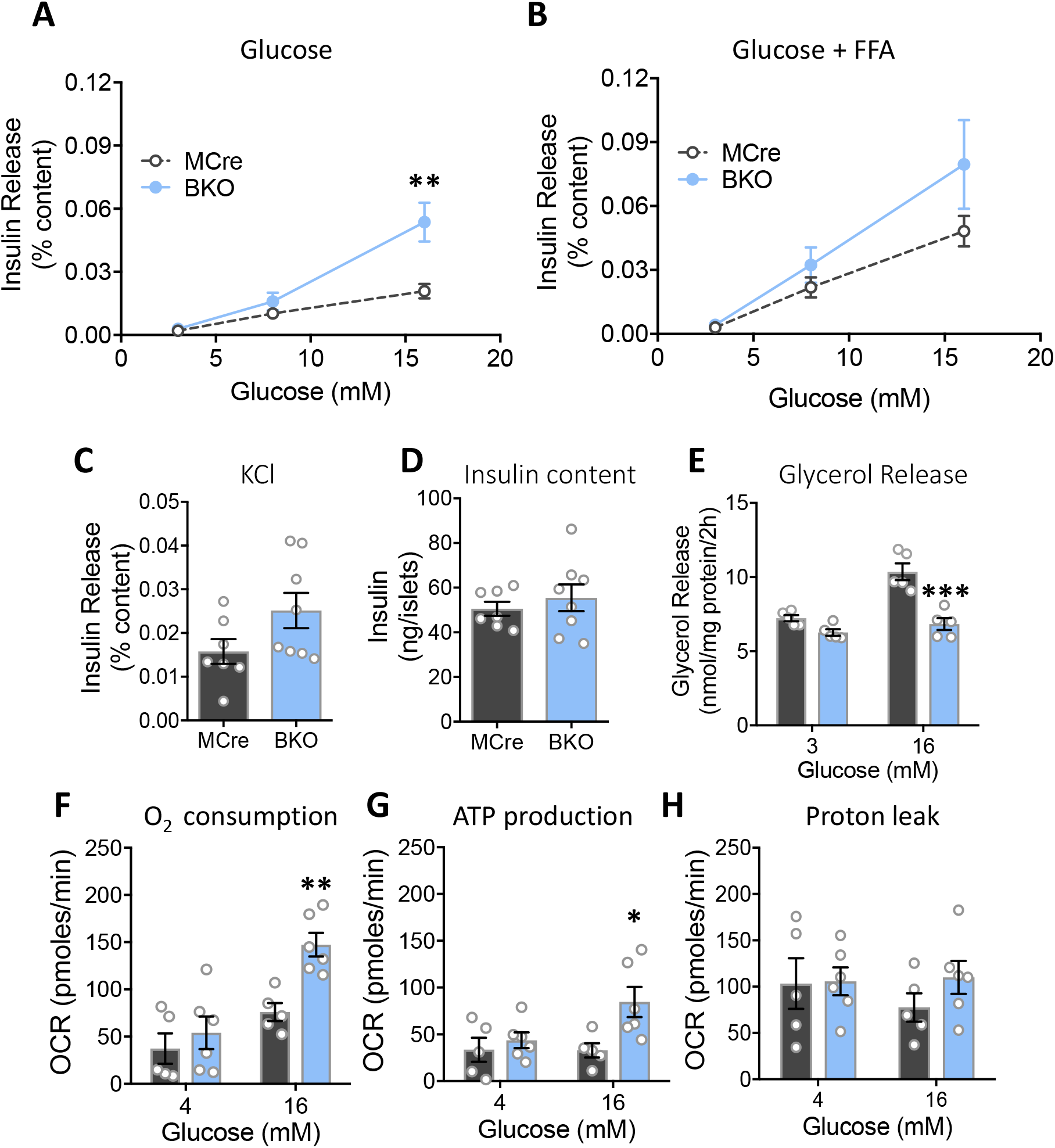
Assessment of *ex vivo* glucose induced insulin secretion and metabolic parameters in BKO versus control mouse islets after 12 weeks of G3PP deletion. (A) Insulin secretion at 3, 8 and 16 mM glucose. (B) Insulin secretion as in A, but in the presence of palmitate/oleate (0.125 mM each). (C) Insulin secretion at 3 mM glucose plus 35 mM KCl. (D) Total insulin content. For A, B, C and D: MCre, n=7; BKO, n=8. (E) Glycerol release (MCre, n=5; BKO, n=5). (F) Oxygen consumption. (G) ATP production. (H) H^+^ leak (MCre, n=5; BKO, n=6). *P < 0.05, **P < 0.01, and ***P < 0.001 vs. MCre (Two-way ANOVA (Panel A and B) and Student’s *t* test (Panel C-H)).

Considering that G3PP, by controlling the availability of Gro3P for mitochondrial electron shuttle, can regulate mitochondrial respiration, we investigated mitochondrial respiration in the BKO islets. Our results showed that in the BKO islets, both oxygen consumption and ATP production were elevated at high glucose concentration but unchanged at low glucose, without affecting H^+^ leak (Figure 3*F*-*H*). The above results indicated that G3PP deletion in β-cells enhances mitochondrial oxidative phosphorylation and GSIS only at high glucose concentration and not at low or physiological basal concentrations.

### Metabolite profiling in G3PP-BKO islets exposed to varying glucose concentrations

To gain insight into the mechanism by which G3PP deletion altered β-cells function, we conducted targeted metabolomics to measure metabolites related to glycolysis, Krebs cycles and additional pathways in BKO and control islets exposed to low, intermediate and high glucose, and correlated the data to GSIS (Figure 4*A*). Results showed that the levels of G3PP substrate Gro3P as well as its precursor dihydroxyacetone phosphate (DHAP) increased with increasing glucose concentration in both BKO and control islets and were markedly higher in the BKO only at high glucose concentration (Figure 4*B* and *C*). This increase in Gro3P and DHAP levels positively correlated with GSIS in both BKO and MCre islets (Figure 4*A*). On the other hand, neither G3PP deletion in β-cells nor glucose concentration had any effect on the islet levels of another known substrate for G3PP, 2-phosphoglycolate (2-PG), which was very low (Figure 4*D*).

**Figure 4.**
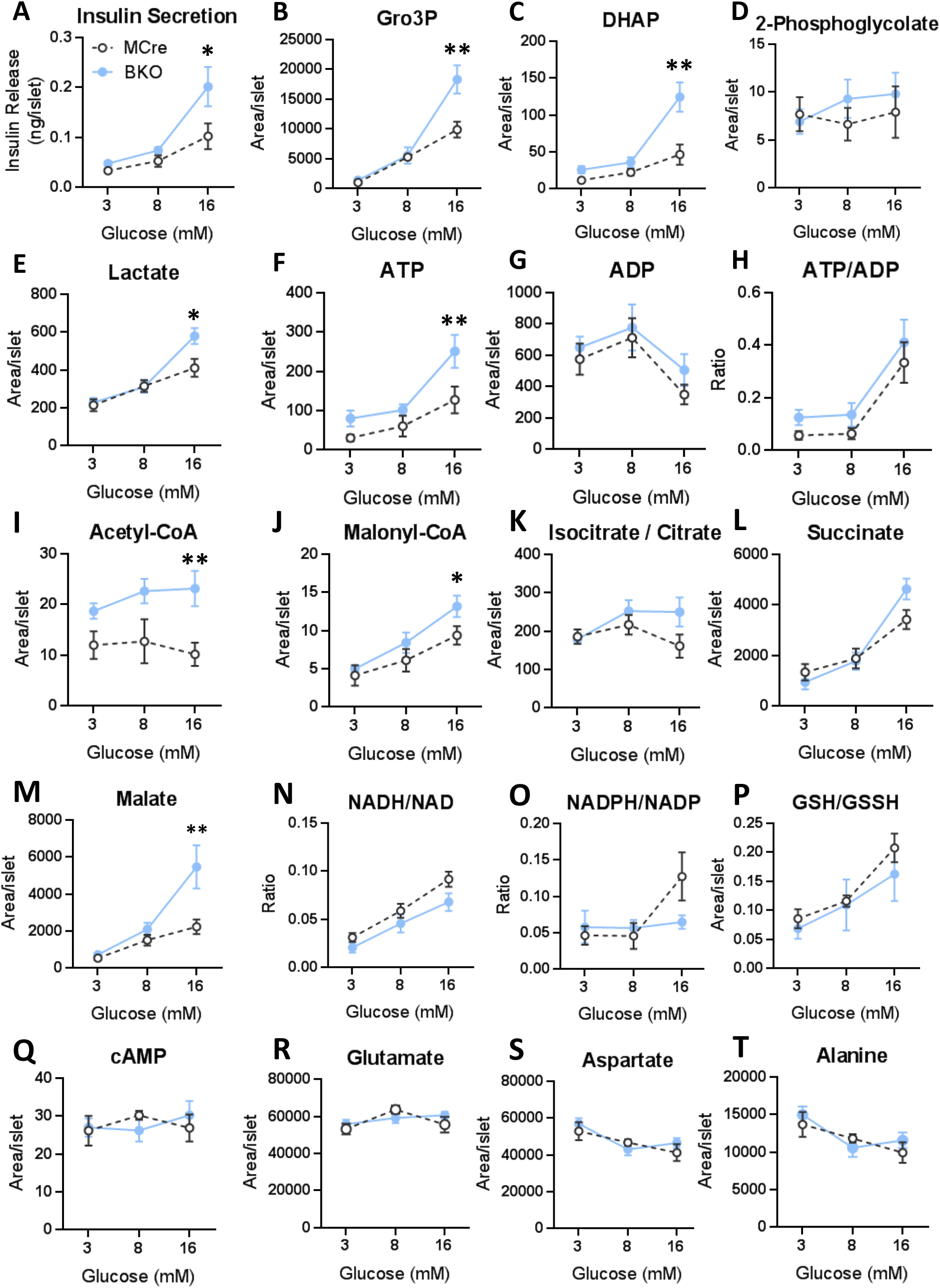
Targeted metabolomics analyses in BKO and control islets, 12 weeks after G3PP deletion. (A) Insulin secretion at 3, 8 and 16 mM glucose after 1h incubation for the metabolomics experiments. At the end of incubation, metabolites were extracted and analyzed by LC-MS/MS. (B) Gro3P. (C) Dihydroxyacetone-phosphate (DHAP). (D) 2-Phosphoglycolate. (E) Lactate. (F) ATP. (G) ADP. (H) ATP/ADP. (I) Acetyl-CoA. (J) Malonyl-CoA. (K) Isocitrate /citrate. (L) Succinate. (M) Malate. (N) NADH/NAD. (O) NADPH/NADP. (P) GSH/GSSH. (Q) cAMP. (R) Glutamate. (S) Aspartate. (T) Alanine. (U) Leucine. Islets from 3 mice were pooled for one measurement and there were 5 such measurements for MCre and BKO, separately. A total of 15 mice were used in each group. Means ± SEM (n= 5 for each group). *P < 0.05 and **P < 0.01 vs. MCre (Two-way ANOVA).

Lactate, an indicator of glycolytic flux, increased with increasing glucose concentration and was higher in BKO islets at 16 mM glucose, compared to the control islets (Figure 4*E*). Similarly, ATP level, an important MCF for insulin secretion, was higher in the BKO islets at 16 mM glucose in association with a small rise in the ATP/ADP ratio (Figure 4*F-H*). BKO islets showed elevated acetyl-CoA levels compared to MCre islets at all glucose concentrations (Figure 4*I*), whereas malonyl-CoA, an important MCF [19] was found to be higher in BKO islets at 16 mM glucose (Figure 4*J*). Interestingly, malonyl-CoA levels also correlated with GSIS results (Figure 4*A*), and implicated the elevated malonyl-CoA, together with ATP, as a plausible metabolites and associated pathway responsible for the elevated GSIS in the BKO islets. Among the Krebs cycle intermediates measured, only malate showed an increase in the BKO islets at 16 mM glucose, whereas no major changes were noticed in the levels of isocitrate/citrate and succinate compared to the control islets (Figure 4*K-M*). There were no differences in AMP, adenosine, GMP, NADH, NAD, NADPH, NADP, GSH, GSSG, fumarate and arginine (Supplemental Figure 3) and in NADH/NAD, NADPH/NADP, GSH/GSSH, cAMP and amino acid levels (Figure 4*N-U*) between the groups. These results showed that G3PP deletion led to increased glycolytic flux in the β-cell, accompanied by elevated acetyl-CoA and malonyl-CoA production and mitochondrial ATP formation, which are known contributors to enhanced GSIS.

### G3PP-BKO islets are more susceptible to chronic glucotoxicity

All the above *in-vivo* and *ex-vivo* metabolic measurements in the BKO mice and islets revealed that G3PP activity is more relevant in the ß-cell at high glucose concentration. Therefore, we examined if G3PP activity impacts the toxic effect of chronic exposure of islets to high glucose levels (glucotoxicity). After incubation of BKO and MCre islets for 7 days at 11 mM glucose (close to fed glucose concentration in mice) and 30 mM glucose (glucotoxicity condition), the BKO islets exposed to 30 mM glucose looked more transparent compared to the control islets (Figure 5*A*). The BKO islets exposed to high glucose showed a significant decrease in insulin content after 7 days incubation (Figure 5*B*). Furthermore, BKO islets showed a marked increase in apoptosis in comparison to control islets after 7 days exposure to glucotoxic conditions (Figure 5*C*). In order to better understand the mechanisms underlying the changes in BKO islets, we measured the expression of different genes related to insulin, ß-cell identity and ER stress. The results revealed that the BKO islets showed a significant decrease in the expression levels of *Ins-2* and *Pdx-1* under glucotoxic conditions compared to control islets (Figure 5*D* and *E*). Expression of *Mafa* showed a more pronounced decrease close to significance (p = 0.056) than *Ins-2* and *Pdx-1* in the BKO islets (Figure 5*F*). There were no differences in these genes, insulin content or apoptosis in WT versus fl/fl mice (Supplemental Figure 5*A-E*). In addition, genes related to glucotoxicity (*Txnip*) and ER stress (*Bip*) showed significantly increased expression under glucotoxicity conditions (Figure 5*G* and *H*; Supplementary Figure 5*F* and *G*) but no difference between groups.

**Figure 5.**
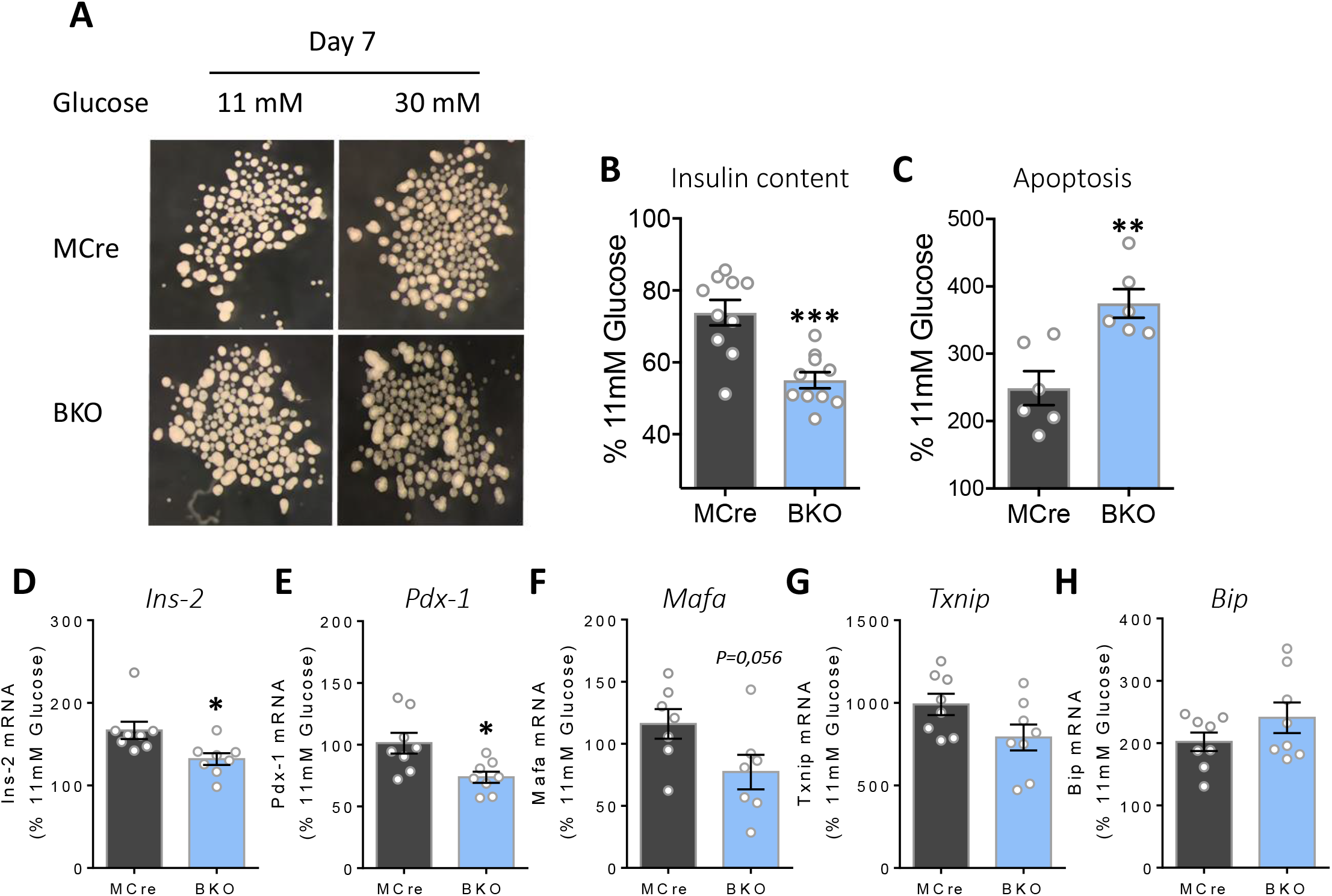
Assessment of chronic glucotoxicity in BKO and control islets. (A) Representative image of islets (Islets from 2 mice were pooled for one measurement) after incubation with 11 and 30 mM glucose for 7 days. After the 7-days incubation, islets were collected to measure: (B) insulin content (MCre, n=10 from 20 mice; BKO, n=10 from 20 mice), (C) apoptosis (MCre, n=6 from 12 mice; BKO, n=6 from 12 mice) and (D-H) gene expression (mRNA) by rt-PCR (MCre, n=8 from 16 mice; BKO, n=8 from 16 mice). (D) *Ins-2*, (E), *Pdx-1* (F) *Mafa*, (G) *Txnip* and (H) *Bip*. Data are presented as % of 11mM glucose from the data shown in Fig 5S. Means ± SEM. *P < 0.05, **P < 0.01, and ***P < 0.001 vs. MCre (Student’s *t* test).

## Discussion

Glucose is the primary nutrient for β-cell stimulation and its metabolism via glycolysis plays a critical role in the regulation of insulin secretion, in part through the production of Gro3P, a metabolite that links glycolysis to GL/FFA cycling, the Gro3P shuttle and direct glycerol release by G3PP [3-5, 11]. However, chronic exposure to excess glucose is toxic to β-cell function [15], and our earlier *in vitro* studies largely done in the tumoral ß-cell line INS1-832/13 suggested that G3PP in β-cells helps in the elimination of excess glucose carbons in the form of glycerol [10]. However, we do not know the glucose concentration range in which the flux from glucose to glycerol via G3PP and Gro3P hydrolysis is most active, independently of lipolysis. The question is important as G3PP shunts glucose carbons from glycolysis into glycerol that escapes the ß-cell. This is an ideal detoxification pathway of excess glucose because glucose carbons leave the cell early-on in their metabolism following glucose entry and its trapping in the form of glucose-6-phosphate by glucokinase, and also because glycerol is a neutral polar molecule that can reach high levels without toxicity [20]. We propose to name this pathway the *“glycerol shunt”*, a new name for a pathway of intermediary metabolism in mammalians whatever is the cell type. It may be highly relevant for cardiometabolic disorders and healthy aging at large as we recently demonstrated, using the nematode *C. elegans*, that G3PP plays key role in the defense against various stresses, including glucotoxicity [21]. In *C. elegans* a mild enhancement in G3PP activity mimics the beneficial effects of calorie restriction, without affecting food intake and fertility [21]. Moreover, a recent human genetic study identified *PGP*/G3PP genetic variants to be associated with longevity in centenarians, together with variants in another gene closely related to glucose metabolism, *FN3KRP* coding for fructosamine-3-kinase related protein [22]. We now provide *in vivo* and *ex vivo* evidence indicating that G3PP controls glucose metabolism, insulin secretion and glucotoxicity in normal β-cells, using an inducible β-cell specific G3PP-KO mouse model.

The results show that loss of G3PP activity in the β-cells in adult stage leads to hyperinsulinemia, even when the mice are fed chow diet and their fed glycemia is comparable to control groups (WT, MCre and fl/fl). This sustained elevated plasma insulin levels in the BKO mice can explain the increased anabolism and energy storage, particularly in the fat, and also the elevated body weight gain, which are not due to altered food intake. The reduced plasma TG levels and increased TG content in visceral fat and skeletal muscle in the BKO mice also suggest that fat is sequestered in peripheral tissues due to the enhanced insulin levels. Thus, insulin activates lipoprotein lipase and promotes fat esterification and inhibits adipose lipolysis, particularly during fasting period [23]. The possibility that the mild glucose intolerance in the BKO mice is due to the increased TG storage in fat and muscle tissues needs to be ascertained. Overall these results suggest that compromised activity of G3PP in β-cells can lead to hyperinsulinemia-driven obesity.

We further verified that the *in vivo* elevated insulinemia in BKO mice is due to intrinsic β-cell effect. Thus, *ex vivo* insulin secretion in isolated islets from the BKO mice at three glucose concentration corresponding to hypoglycemia, normoglycemia and hyperglycemia, 3, 8, and 16 mM respectively, is elevated only at the highest concentration of glucose. This increase was associated with a marked decline in glycerol production and increase in cellular levels of Gro3P, which is the starting substrate for GL/FFA cycle. Thus, the flux through the GL/FFA cycle, particularly through its lipogenic segment, is expected to increase in BKO islets, leading to the production of MCF (e.g., diacylglycerol and monoacylglycerol) necessary for the lipid amplification of GSIS [5]. Moreover, the high glucose concentration dependency for the elevated insulin secretion is not masked by the presence of FFA, suggesting that Gro3P accumulation is the driving force for the enhanced GSIS in BKO islets. The present findings emphasize that G3PP deletion in β-cells has no significant impact at low and intermediate/physiological glucose concentration (3 and 8 mM) on GSIS and on the glycolytic and Krebs cycle metabolites, compared to the control islets. Only at 16 mM glucose there is a significant increase in Gro3P and DHAP levels in BKO islets. It is likely that at intermediate glucose concentration, β-cells use Gro3P effectively for the Gro3P shuttle, lipogenesis and lower-glycolysis, without much build-up of this substrate. Furthermore, deletion of G3PP also led to elevated glycolysis in the islets, specifically at high glucose concentration, probably due to increased diversion of Gro3P towards lower glycolysis and lactate production. G3PP deficient islets displayed increased levels of some Krebs cycle intermediates and mitochondrial respiration only at high glucose concentration, as evidenced by elevated O_2_ consumption and ATP production, likely because of augmented pyruvate production and Gro3P shuttle mediated electron supply to mitochondrial electron transport chain, coupled with oxidative phosphorylation. Thus, it is relevant to note that the NADH/NAD levels in BKO islets are lower at 16 mM glucose, without any change in NADH levels, suggesting an efficient transfer of reducing equivalents from cytosol to mitochondria via the Gro3P shuttle. The increased ATP availability also can explain, in part, enhanced GSIS in the BKO islets through the closure of K_ATP_ channels [3].

Accelerated GSIS amplification through the acetyl-CoA carboxylase (ACC)/malonyl-CoA (Mal-CoA)/ carnitine palmitoyltransferase 1 (CPT-1) signaling network [3, 24, 25] is a likely possibility in the BKO islets as there is a marked increase in the levels of acetyl-CoA, the substrate for ACC, and malonyl-CoA, produced by ACC, under high glucose levels. Overall, the results indicate that G3PP deletion in β-cell activates multiple pathways including ATP production, the GL/FFA cycle and ACC/malonyl-CoA/CPT-1 network to achieve augmented insulin secretion at high glucose concentrations and these effects are related to reduced activity of the glycerol shunt and elevated Gro3P levels (Figure 6).

**Figure 6.**
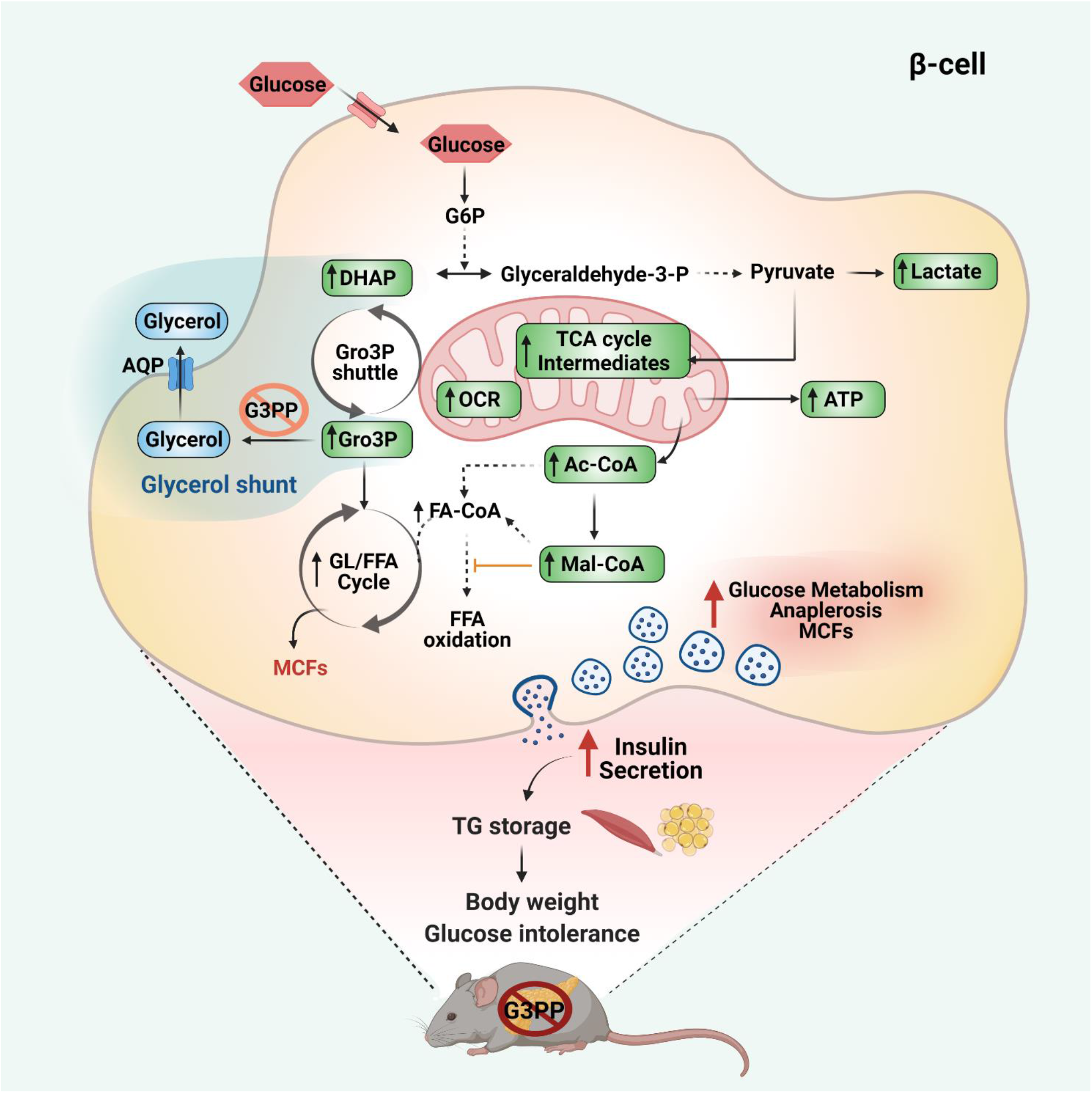
Model depicting the effect of G3PP ß-cell specific deletion on energy homeostasis and body weight via increasing insulin secretion under high glucose concentration. The abbreviations are: Ac-CoA, acetyl-CoA; AQP, aquaporin channel; ATP, adenosine triphosphate; DHAP, dihydroxyacetone phosphate; FA-CoA, fatty acyl-CoA; FFA, free fatty acid; G3PP, glycerol 3-phosphate phosphatase; G6P, glucose 6-phosphate; GL/FFA cycle, glycerolipid/ free fatty acid cycle; Glyceraldehyde-3-P, glyceraldehyde-3-phosphate; Gro3P, glycerol 3-phosphate; Mal-CoA, malonyl-CoA; MCF, metabolic coupling factor; OCR, oxygen consumption rate; TCA cycle, tricarboxylic acid cycle; TG, triglyceride.

The results also emphasized that Gro3P is the more plausible physiological substrate for G3PP in the β-cell as 2-PG levels, the other suggested G3PP substrate, were found to be very low and did not change in control and BKO islets at all tested glucose concentrations [11, 13]. Earlier an *in vitro* studies in cancer cell lines suggested that G3PP/PGP may act as a ‘metabolite repair enzyme’ by eliminating toxic by-products of glycolysis, 4-phosphoerythronate and 2-phospho-L-lactate [26]. However, we did not notice any toxicity or dysfunction of G3PP-deleted β-cells at physiological glucose concentration, both *in vivo* and *ex vivo*. Instead, these ß-cells displayed more efficient metabolic activation and insulin secretion, showing indirectly that under normal conditions, the above toxic metabolites do not accumulate or cause toxicity in the BKO ß-cells [11, 13].

As the studies above indicated that lack of G3PP in β-cells impacts metabolism and insulin secretion only at elevated glucose concentrations, we found it to be of importance to determine if this enzyme plays a role in preventing ß-cell glucotoxicity and nutri-stress [15], a term encompassing the toxic actions of all nutrients in excess: carbohydrates, lipids and proteins, reflected by elevated blood levels of glucose, TG and FFA as well as some amino acids. Chronic exposure of BKO isolated islets to 30 mM glucose increased apoptosis compared to MCre islets and reduced insulin content and the expression of *Ins1, Pdx1* and *Mafa*, that are crucial for β-cell function [15, 27, 28]. Reduction in the expression of these genes is known to induce dedifferentiation, a feature of β-cells dysfunction [15, 28]. These findings indicate that G3PP in the ß-cell plays a role in glucodetoxification, maintaining the differentiated state and the expression of the insulin gene, *Ins1*.

In conclusion, the data demonstrate that the *PGP* gene product encodes for a protein that shows Gro3P phosphatase activity and that deletion of this gene specifically in pancreatic β-cells has profound effects on glucose metabolism, β-cell function and GSIS, only at high glucose concentration (Figure 6). The data identify a novel metabolic pathway, the *glycerol shunt*, implicated in glucodetoxification and fuel partitioning via the metabolic fate of Gro3P and DHAP. We propose that G3PP in β-cells acts as a security valve that expels excessive glucose carbons as glycerol to prevent more than necessary secretion of insulin that could cause obesity and possibly hypoglycemia, as well as ß-cell dysfunction and death under chronic hyperglycemic conditions. The findings are potentially of general interest as we know very little about fuel excess detoxification pathways at large, whatever is the cell type, and because of the recently emerging view that G3PP is implicated in healthy aging [21, 22].

## Research Design and Methods

### Generation of G3PP conditional KO mice and breeding strategy

Heterozygous G3PP-lox/lox mice in which exons 1 and 2 of *PGP* (coding for G3PP protein) gene are flanked with LoxP sites were generated by Ingenious Targeting Laboratory (Stony Brook, NY). The donor targeting vector was designed so that the first loxP sequence is inserted on the 5’ side of exon 1 and the second at the 3’ end of the FRT-flanked Neo selection cassette (after exon 2) (Figure 1*A*). The targeting vector was then transfected by electroporation to FLP C57BL/6 ES cells. Positive targeted ES clones, where the Neo cassette was removed by the FLP recombinase, were microinjected into Balb/c blastocysts to generate chimeras. These chimeras were crossed to wildtype (WT) C57Bl/6N to generate heterozygous G3PP-lox/+ mice. Heterozygous G3PP-lox/+ mice were bred to produce homozygous G3PP-lox/lox (G3PP^flox/flox^) mice. Homozygous G3PP-lox/lox were then crossed with heterozygous Mip-CreERT2 mice [17] to obtain Mip-CreERT2/+, G3PP-flox/+ and G3PP-flox/+ mice. These mice were further mated to get MipCre-ERT2/+, WT, G3PP-flox/flox and Mip-CreERT2/+; G3PP-flox/flox mice. The presence of the WT or the floxed *PGP* allele was evaluated in DNA from ear punch tissue fragments using the following primers: NDEL1: 5’-ACCTCTTGCCTGCCCTTCAAGG-3’; NDEL2: 5’-TTGACCCATTTCAGTCTCAGCAACAGG-3’, and the presence of Cre transgene using the following primers: forward: 5’-CCTGGCGATCCCTGAACATGTCCT-3’, Reverse: 5’-TGGACTATAAAGCTGGTGGGCAT -3’. Specific amplification of a 452 bp DNA fragment corresponding to the floxed-*PGP* allele, a 312 bp fragment corresponding to the WT *PGP* allele and the presence of the Mip-Cre transgene by a 267 bp DNA fragment was verified by genomic PCR (Figure 1*B*).

### Animals

All mice were on C57BL/6N genetic background. At 8 weeks of age, wild-type (WT), G3PP^flox/flox^ (fl/fl), Mip-CreERT2 (MCre) and Mip-CreERT2/^+^;G3PP^flox/flox^ male mice received tamoxifen (TMX) intraperitoneal injections for 5 consecutive days (50 mg/kg body weight, dissolved in 90% corn oil plus 10% ethanol) to induce Cre recombinase and β-cell specific deletion of G3PP in the Mip-CreERT2/^+^; G3PP ^flox/flox^ mice (BKO). After 5 days of TMX injections, mice were placed in individual cages and fed normal chow diet (15% fat by energy; Harlan Teklad, Madison, WI, USA) for 12 weeks. Body weight and food intake were measured weekly. Specific deletion of G3PP expression in pancreatic islets without any change in other tissues was ascertained in the BKO mice, as compared to WT, MCre and fl/fl control mice (Figure 1*C*). Body composition (lean and fat mass) after 12 weeks after TMX injection was measured by magnetic resonance imaging (EchoMRI Analyzer-700). Animals were housed individually at room temperature (23°C) with a 12 h light/12 h dark cycle. 20-week-old mice were anesthetized by intraperitoneal injection of ketamine (100 mg/kg)/xylazine (10 mg/kg), followed by cardiac puncture and euthanasia via cervical dislocation. Various tissues, including adipose depots, liver, kidney, etc., were immediately collected and kept frozen for further analyses. All procedures were approved by the institutional committee for the protection of animals (Comité Institutionnel de Protection des Animaux du Centre Hospitalier de l’Université de Montréal).

### Pancreatic islet isolation

Pancreatic islets were isolated from BKO, WT, MCre and fl/fl mice as previously described [29]. After isolation, the islets were kept in RPMI medium supplemented with 2 mM glutamine, 1 mM sodium pyruvate, 10 mM HEPES (pH 7.4), penicillin (100U/ml)/ streptomycin (100μg/ml), 10% FBS and containing 5 mM glucose (glucose concentration during overnight recovery was 8 mM for glucotoxicity experiments) for overnight recovery and then employed for *ex-vivo* insulin secretion experiments and other measurements as detailed below.

### Western blotting

Mouse tissues and pancreatic islets were lysed in RIPA buffer (10 mM Tris-HCl, 1 mM EDTA, 0.5 mM EGTA, 1% Triton X-100, 0.1% sodium deoxycholate, 0.1% sodium dodecylsulfate and 140 mM NaCl) and the protein extracts were used for western blot analysis to validate β-cell specific G3PP deletion (G3PP primary antibody: PGP antibody E-10, sc-390883; secondary antibody: m-IgGκ BP-HRP, sc-516102). For all tissues, α-tubulin was used as the gel loading control.

### Intraperitoneal glucose tolerance test (IPGTT)

IPGTT was performed on 16-week-old male G3PP-BKO, WT, MCre and fl/fl mice. Glucose (2 g/kg body weight) was injected intraperitoneally in conscious mice at 13:00 hours after 6 h of food withdrawal. Glycemia was measured in the tail blood at time 0, 15, 30 and 60 min following glucose administration using a glucometer (Contour blood glucometer, Bayer). Insulin levels were also measured in the tail blood.

### Insulin tolerance test (ITT)

ITT was performed on 19-week-old male G3PP-BKO, WT, MCre and fl/fl mice. Insulin (0.75 U/kg body weight, Humulin; Lilly, Indianapolis, USA) was injected intraperitoneally (IPITT) in conscious mice at 13:00 hours, after 4h of food withdrawal. Glycemia was measured in the tail blood at time 0, 15, 30, 45, 60, 90 and 120 min using glucometer. Blood was collected from the tail at time 0 min to measure plasma insulin levels.

### Plasma parameters

Twenty week old mice fed a normal diet were anesthetized as described above and blood was collected through cardiac puncture at 08:00 hour. Plasma glucose, insulin, non-esterified fatty acids (NEFA), TG and glycerol were measured [30].

### *Ex vivo* insulin secretion

After overnight recovery, islets were transferred in complete RPMI medium containing 3 mM glucose for 2h and preincubated for 45 min at 37°C in Krebs Ringer buffer with 10 mM Hepes, pH 7.4 (KRBH) containing 0.5% defatted-BSA, 2 mM glutamine, 50 μM L-carnitine and 3 mM glucose. Batches of 10 islets, in triplicates, were incubated for 1h at 3, 8 and 16 mM glucose in the presence or absence of palmitate/oleate (0.15 mM/each, complexed with BSA) and at 3 mM glucose plus 35 mM KCl. Insulin release was normalized for the total islet insulin content.

### Glycerol release

Batches of 100 islets that were preincubated for 1h in KRBH containing 0.5% defatted-BSA, 2 mM glutamine), 50 μM L-carnitine and 4 mM glucose. After preincubation, the islets were incubated for 2h at 4 and 16 mM glucose. Glycerol release into the medium was determined using [γ-^32^P]ATP (Perkin Elmer Life Sciences) and glycerol kinase (Sigma) [31].

### Oxygen consumption and mitochondrial function

Isolated mouse islets (batches of 75 islets) were transferred to XF24 islet capture microplates in 4 mM KRBH (0.07% defatted-BSA, 2 mM glutamine and 50 μM L-carnitine). After basal respiration measurement in a XF24 respirometer (Seahorse Bioscience) for 20 min, glucose levels were increased to 16 mM and O_2_ consumption was measured for another 1h. Then oligomycin (to assess uncoupled respiration), FCCP (to estimate maximal respiration), and antimycin/rotenone (to measure non-mitochondrial respiration) were added successively.

### Glucotoxicity

After overnight recovery in RPMI medium containing 8 mM glucose and 10% FBS, 150 -200 islets were incubated for 7 days at 11 and 30 mM glucose in serum-free RPMI medium with 0.5% BSA. Media were changed on alternate days. After 7 days incubation, islets were collected to measure insulin content, gene expression and apoptosis (Cell Death Detection kit, Roche, Basel, Switzerland) [32].

### RNA extraction and quantitative PCR

Total RNA was extracted from 150 islets (RNeasy Micro Kit, Qiagen). Following reverse-transcription to cDNA, expression of different genes was determined by quantitative PCR using SYBR Green (Qiagen (QuantiTect)). All gene expression analyses were conducted in duplicate and normalized to the expression of *18S*. Sequences for the primers used are listed in Supplementary Table 1.

### Targeted metabolomics

Isolated islets (250) were incubated in 1.5 ml tubes under similar conditions as for the *ex vivo* insulin secretion experiments. After incubation, media were collected to measure insulin secretion and 675 μl of ice-cold extraction buffer (80% methanol, 2 mM ammonium acetate, pH 9.0) was quickly added to the islets and mixed by vortex, followed by sonication in a cup-horn (Q700 Sonicator, Qsonica, Newtown, CT) at 150 watts for 1 min (cycles of 10s on, 10s off) in an ethanol-ice bath. Islet extracts were centrifuged at 4°C for 10 min at 20,000×g, and 600 μl supernatant from each condition was collected in separate ice-cold glass tubes to which water (168 μl) was added. Polar metabolites were extracted with 960 μl of chloroform:heptane (3:1, v/v) by 2 × 10s vigorous mixing by vortex followed by 10-min incubation on ice and 15 min centrifugation at 4°C at 4,500×g. The upper aqueous phase (540 μl) was transferred in a cold 1.5 ml tube, followed by centrifugation and the supernatants (400 μl) were collected. All the samples were frozen in liquid nitrogen and freeze-dried, and stored at −80°C until used. The dried samples were reconstituted in 20 μl of Milli-Q water at 4°C, and 5 μl injections were employed in duplicate for analysis on an LC-electrospray ionization-MS/MS system (Agilent 1200 SL and a triple-quadrupole mass spectrometer (4000Q TRAP MS/MS, Sciex)) [10]. Peak areas were used for relative quantification of identified metabolites.

## Statistical analyses

Data were analyzed using GraphPad Prism 6 (GraphPad Software, San Diego, CA). Results are expressed as means ± SEM. Statistical differences between two groups were assessed by unpaired, two-tailed Student’s *t* test, and between multiple groups using one-way or two-way analyses of variance (ANOVA) with Bonferroni or Student *t* test, as indicated. A *p* value < 0.05 was considered statistically significant.

## Study approval

All the procedures for mice studies were performed in accordance with the Institutional Committee for the Protection of Animals at the University of Montréal CRCHUM.

## Author contributions

AAM, MLP, SRMM, and MP designed research; AAM, PP, MLP, RL, EL, JG and YM conducted experiments; AAM, MLP, SRMM, and MP analyzed data; PP, MLP, EP, RS, RA and FAM reviewed and edited the manuscript and AAM, SRMM and MP wrote the paper.

## Acknowledgments

This study was supported by funds from Canadian Institutes of Health Research (to M.P. and S.R.M.M.), and from Dasman Diabetes Research Institute/ Montreal Medical International (to M.P, MSRM, RA and FA-M). M.P. was recipient up to 2019 of a Canada Research Chair in Diabetes and Metabolism. AA-M is supported by a scholarship from Kuwait University. PP was recipient of postdoctoral fellowships from the Fondation Valifonds, the Montreal Diabetes Research Center, the CRCHUM and Mitacs. We thank the core facilities for Cellular Physiology, Metabolomics and Rodent Phenotyping of the CRCHUM/Montreal Diabetes Research Center. We thank Jennifer Estall (University of Montreal, Canada), Erik Joly (CRCHUM) and Julien Lamontagne (CRCHUM) for helpful advice and discussion. We also thank Isabelle Chénier and Heidi Erb for technical assistance with animal experiments. Figure 6 was created with BioRender.com.

