## Supplementary material for "A glycerol shunt functions as a glucose excess security valve in pancreatic β-cells"

**Supplemental material**

**Figure S1**. *In vivo* parameters of WT and G3PP-fl/fl mice. (A) Body weight. (B) Percentage of body weight gain. (C) Food intake (For A, B and C: WT, n=7; fl/fl, n=11). (D) Plasma parameters in fed mice (For glycemia: WT, n=9; fl/fl, n=6; for insulin, C-peptide, TG, glycerol and FFA: WT, n=9; fl/fl, n=10). (E) Tissue weights of VAT, SAT, liver, BAT and brain (WT, n=5; fl/fl, n=7; MCre, n=6; BKO, n=7). (F) Glycemia during IPGTT (WT, n=6; fl/fl, n=11); inset depicts area under the curve (AUC) for glycemia after 120 min. (G) Insulinemia during IPGTT (WT, n=6; fl/fl, n=11); inset depicts AUC for insulinemia after 120 min. (H) Glycemia during ITT (WT, n=7; fl/fl, n=9 out of 11 mice as two mice showing stress were removed). Inset depicts area above the curve (AAC) after 120 min. Means ± SEM. *P < 0.05 vs. MCre and #P < 0.05 and ##P < 0.01 vs. WT and fl/fl (Two-way ANOVA (Panel A, C, F, G and H), Student's *t* test (Panel B and D) and One-way ANOVA (Panel E)).

**Figure S2**. *Ex vivo* insulin secretion, glycerol release and $O_2$ consumption in WT and G3PP-fl/fl mouse islets. (A) Insulin secretion at 3, 8 and 16 mM glucose. (B) Insulin secretion as in A, but with added palmitate/oleate (0.125 mM each). (C) Insulin secretion at 3 mM glucose plus 35 mM KCl. (D) Total insulin content (For A, B, C and D: WT, n=6; fl/fl, n=8) (E) Glycerol release (WT, n=5; fl/fl, n=5). (F) $O_2$ consumption. (G) ATP production. (H) $H^+$ leak (For F, G and H: WT, n=6; fl/fl, n=5) (I) Basal respiration. (J) Maximal respiration. (K) Non-mitochondrial respiration (For I, J and K: WT, n=6; fl/fl, n=5; MCre, n=5; BKO, n=6). Means ± SEM. (Two-way ANOVA (Panel A and B), Student's *t* test (Panel C-H) and One-way ANOVA (Panel I-K)).

**Figure S3**. Targeted metabolomics determination in islets incubated at various glucose concentrations. BKO and control mouse islets were exposed to 3, 8 and 16 mM glucose for 1h.

Then, metabolites were extracted and analyzed by LC-MS/MS: (A) ANP, (B) adenosine, (C) GMP, (D) leucine, (E) arginine, (F) GSH, (G) GSSG, (H) fumarate, (I) NADH, (J) NAD, (K) NADPH, (L) NADP. Islets from 3 mice were pooled for one measurement and there were 5 such measurements for MCre and BKO, separately. A total of 15 mice were used in each group. Means ± SEM (n= 5 for each group) (Two-way ANOVA).

**Figure S4**. Glucotoxicity in WT and G3PP-fl/fl mouse islets. (A) Representative photo of islets incubated at 11 and 30 mM glucose for 7 days. Islets incubated as In A (Islets from 2 mice were pooled for one measurement) were collected to measure: (B) insulin content (WT, n=6 from 12 mice; fl/fl, n=7 from 14 mice), (C) apoptosis and (D-H) gene expression (WT, n=5 from 10 mice; fl/fl, n=5 from 10 mice) (mRNA) by rt-PCR: (D) *Ins-2*, (E), *Pdx-1* (F) *Mafa*, (G) *Txnip* and (H) *Bip*. Data are presented as % of 11mM glucose from the data shown in Figure 5S; Means ± SEM (Student's *t* test).

**Figure S5**. Glucotoxicity in BKO islets and control mouse islets. Islets (Islets from 2 mice were pooled for one measurement) were incubated at 11 and 30 mM glucose for 7 days and were collected to measure: (A) Insulin content (WT, n=7; fl/fl, n=7; MCre, n=8; BKO, n=8). (B) Apoptosis (WT, n=6; fl/fl, n=7; MCre, n=10; BKO, n=10). (C-G) Gene expression (WT, n=5; fl/fl, n=5; MCre, n=6; BKO, n=6), mRNA measured by rt-PCR: (C) *Ins-2*, (D), *Pdx-1* (E) *Mafa*, (F) *Txnip* and (G) *Bip*. Means ± SEM. *P < 0.05, **P < 0.01, ***P < 0.001 and ****P < 0.0001 vs. 11mM of the same genotype and #P < 0.05, ##P < 0.01, ###P < 0.001 and ####P < 0.0001 vs. 30mM WT and $P < 0.05 and $$P < 0.01 vs. 30mM fl/fl and &P < 0.05 vs. 30mM MCre (Two-way ANOVA).

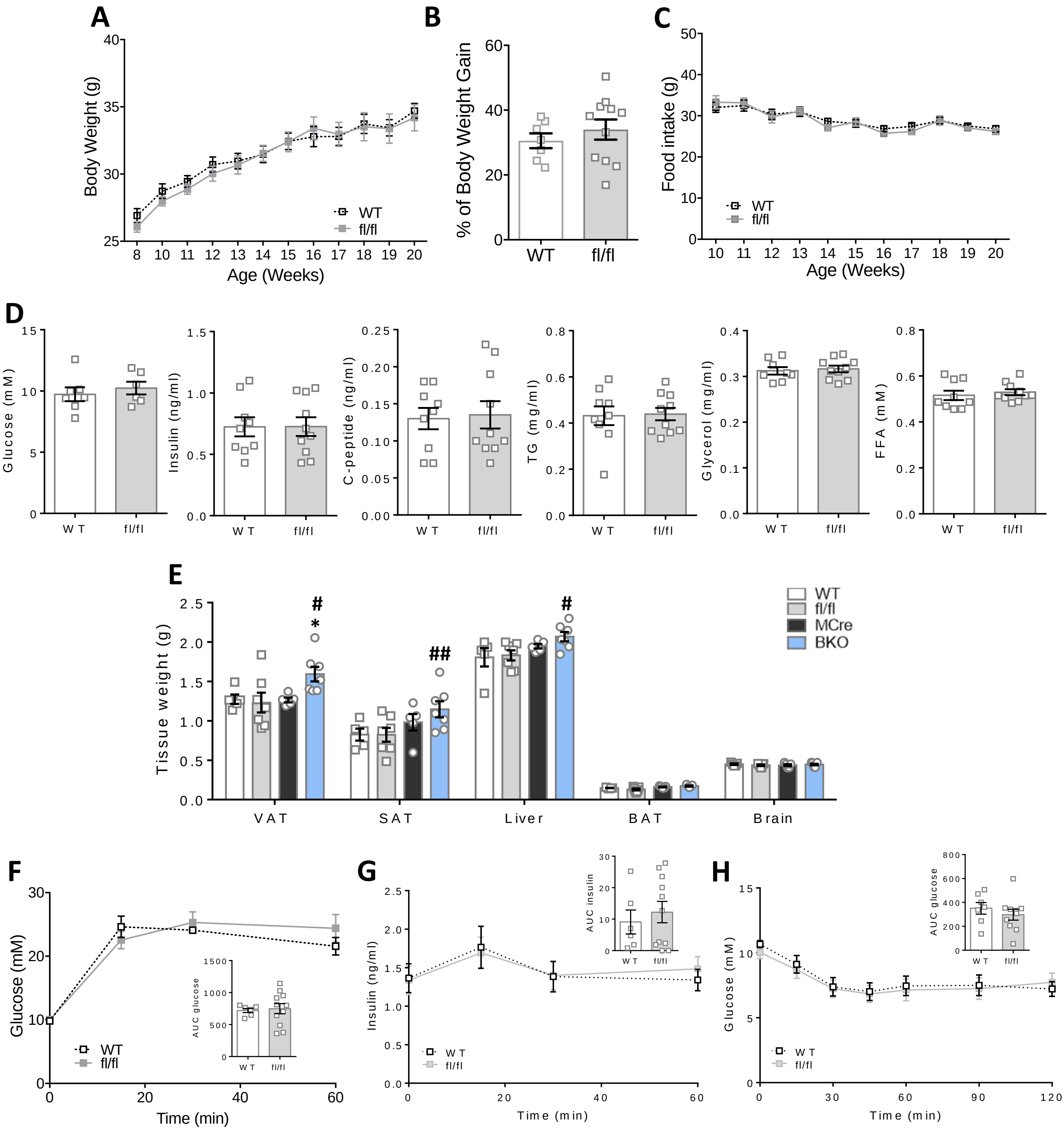
A
Body Weight (g)
Age (Weeks)
WT
fl/fl
B
% of Body Weight Gain
WT
fl/fl
C
Food intake (g)
Age (Weeks)
WT
fl/fl
D
Glucose (mM)
Insulin (ng/ml)
C-peptide (ng/ml)
TG (mg/ml)
Glycerol (mg/ml)
FFA (mM)
WT
fl/fl
E
Tissue weight (g)
WT
fl/fl
MCre
BKO
VAT
SAT
Liver
BAT
Brain
F
Glucose (mM)
Time (min)
AUC glucose
G
Insulin (ng/ml)
Time (min)
AUC insulin
H
Glucose (mM)
Time (min)
AUC glucose

Figure S1

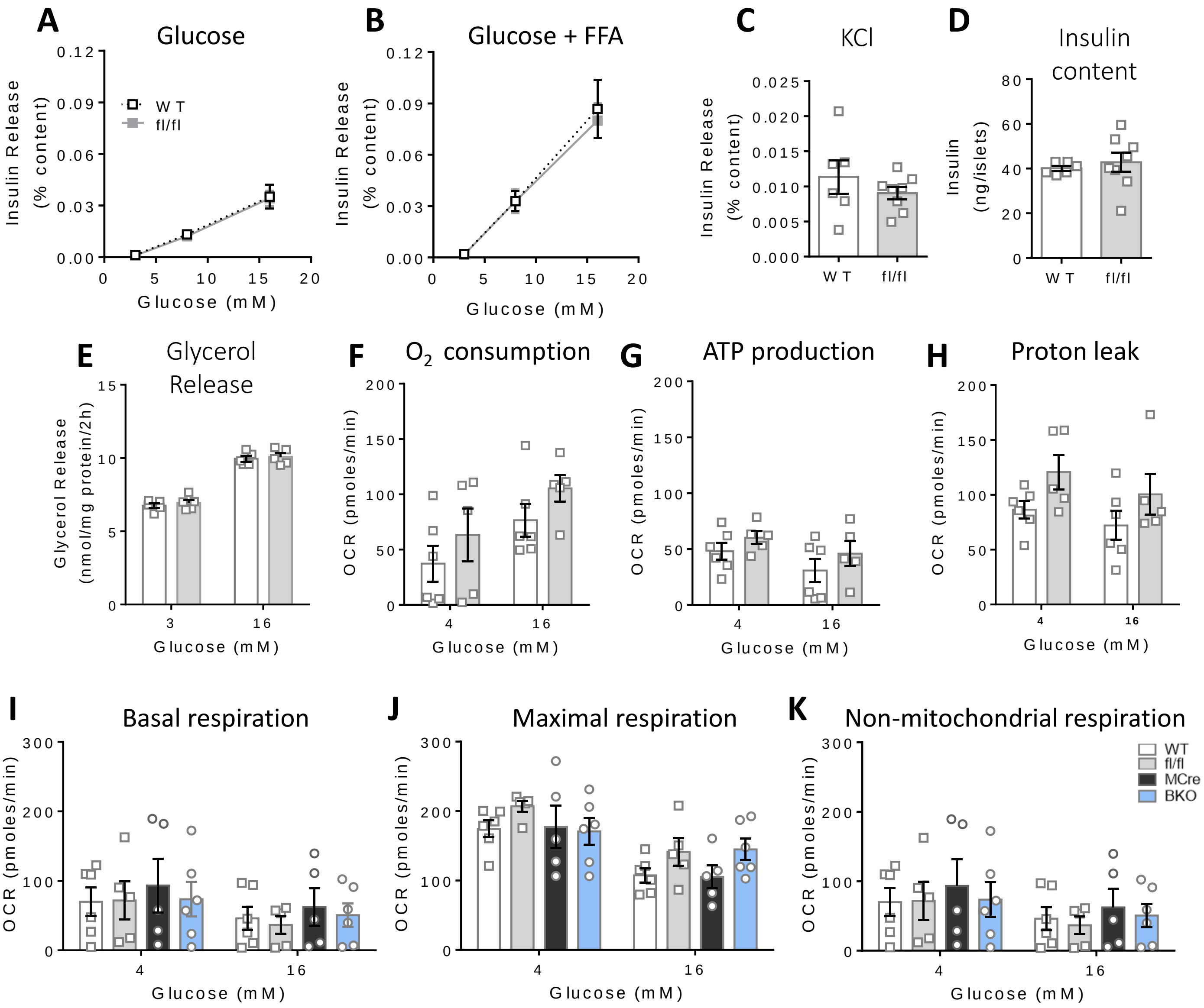
A
Glucose
Insulin Release (% content)
W T
fl/fl
Glucose (mM)
B
Glucose + FFA
C
KCl
D
Insulin content
Insulin (ng/islets)
E
Glycerol Release
Glycerol Release (nmol/mg protein/2h)
F
O2 consumption
OCR (pmoles/min)
G
ATP production
H
Proton leak
I
Basal respiration
J
Maximal respiration
K
Non-mitochondrial respiration
WT
fl/fl
MCre
BKO

Figure S2

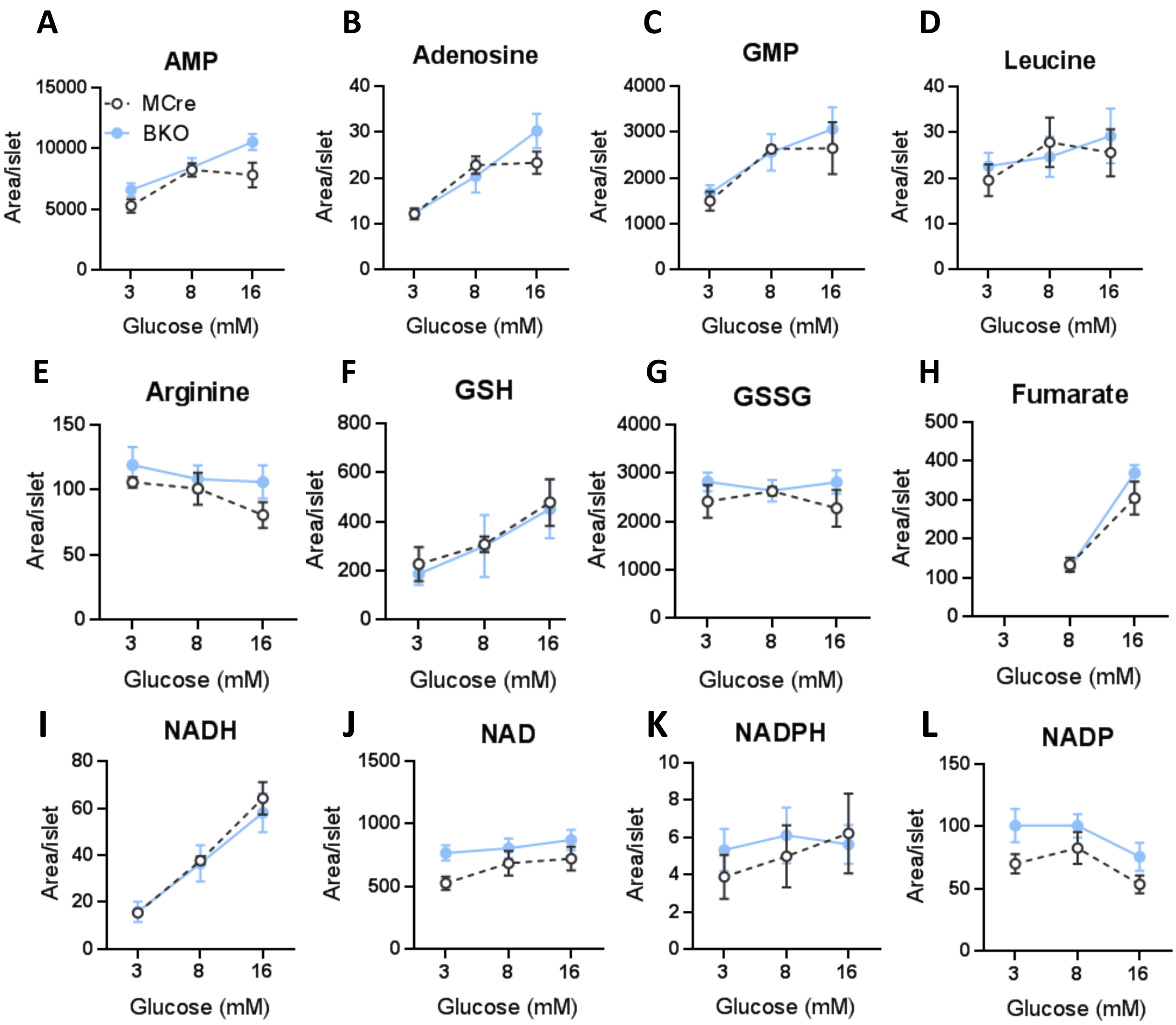
A
AMP
MCre
BKO
Area/islet
Glucose (mM)
B
Adenosine
C
GMP
D
Leucine
E
Arginine
F
GSH
G
GSSG
H
Fumarate
I
NADH
J
NAD
K
NADPH
L
NADP

Figure S3

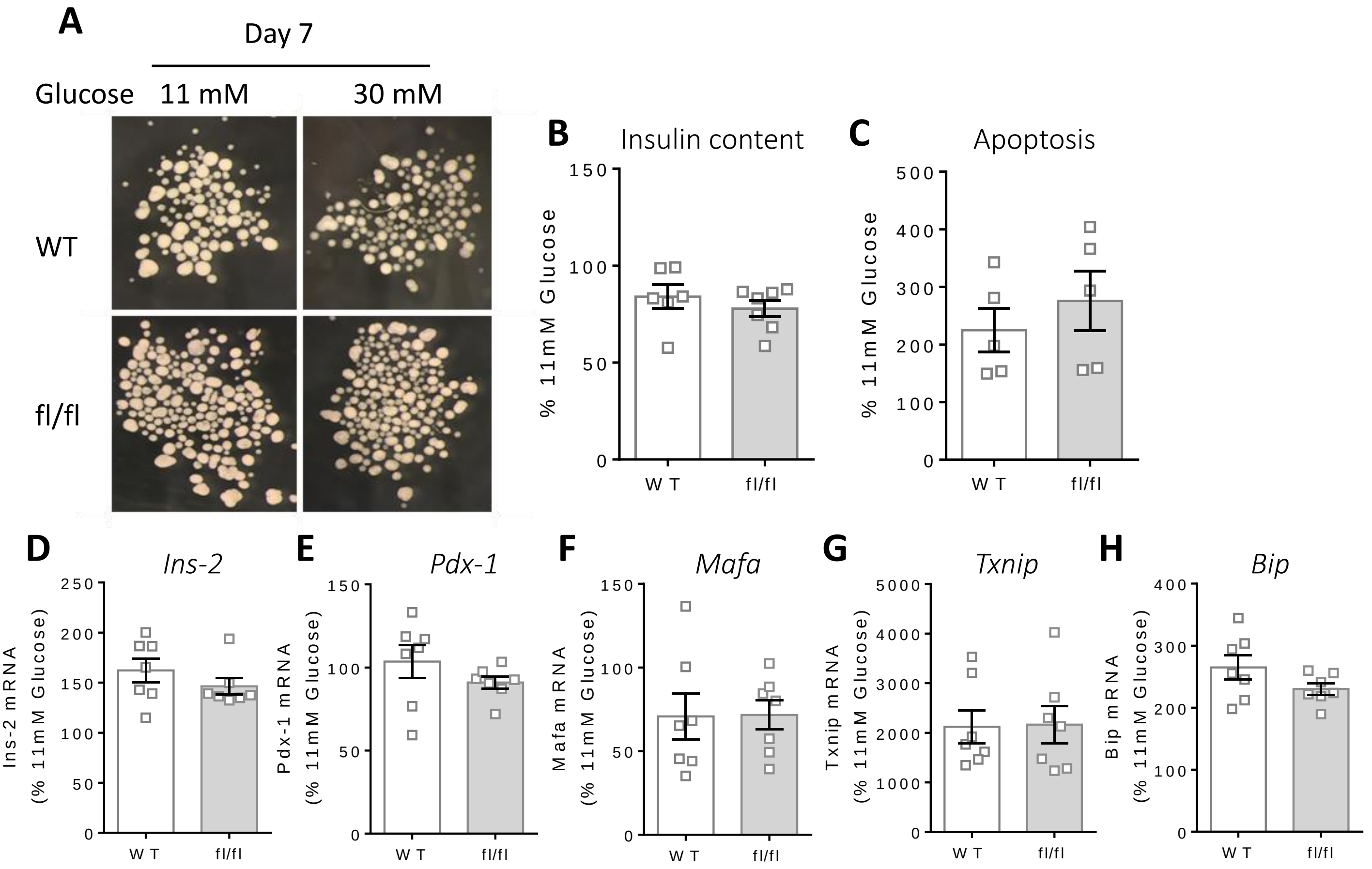
A
Day 7
Glucose 11 mM 30 mM
WT
fl/fl
B Insulin content
% 11mM Glucose
C Apoptosis
% 11mM Glucose
WT fl/fl
D Ins-2
Ins-2 mRNA (% 11mM Glucose)
E Pdx-1
Pdx-1 mRNA (% 11mM Glucose)
F Mafa
Mafa mRNA (% 11mM Glucose)
G Txnip
Txnip mRNA (% 11mM Glucose)
H Bip
Bip mRNA (% 11mM Glucose)

Figure S4

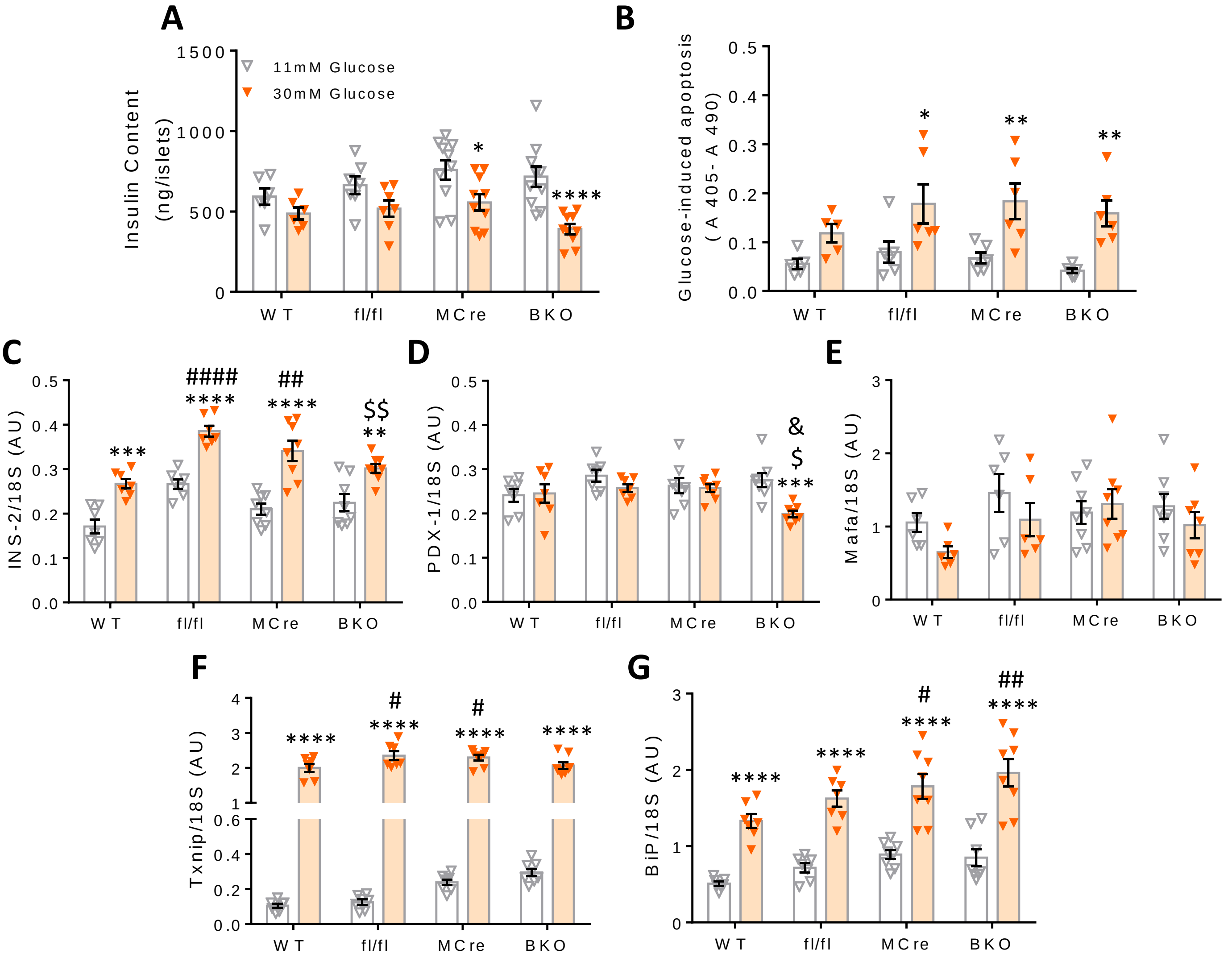
A
B
C
D
E
F
G
11mM Glucose
30mM Glucose
Insulin Content (ng/islets)
Glucose-induced apoptosis (A 405- A 490)
INS-2/18S (AU)
PDX-1/18S (AU)
Mafa/18S (AU)
Txnip/18S (AU)
BiP/18S (AU)
WT
fl/fl
MCre
BKO

Figure S5

Table S1:Primer sequences used for RT-PCR

| Gene | Primer sequences (5′-3′) |
|---|---|
| *Ins-2* | F:TGGAGGCTCTCTACCTGGTG |
| | R:TCTACAATGCCACGCTTCTG |
| *Mafa* | F:GTGCTGGAGGATCTGTACTGG |
| | R:ATGGTGGTGATGGTGATGG |
| *Pdx-1* | F:GGTATAGCCGGAGAGATGC |
| | R:CTGGTCCGTATTGGAACG |
| *Bip* | F:TGCAGCAGGACATCAAGTTC |
| | R:TACGCCTCAGCAGTCTCCTT |
| *Txnip* | F:CGAGTCAAAGCCGTCAGGAT |
| | R:TTCATAGCGCAAGTAGTCCAAAGT |
| *18s* | F:CTG AGA AAC GGC TAG CAC ATC |
| | R:GGC CTC GAA AGA GTC CTG TAT |
